# Combining mutation detection with fragmentomics features leads to improved tumor-informed ctDNA detection

**DOI:** 10.64898/2026.03.30.714025

**Authors:** Yixin Lin, Carmen Oroperv, Amanda Frydendahl Boll Johansen, Mads Heilskov Rasmussen, Claus Lindbjerg Andersen, Søren Besenbacher

## Abstract

Liquid biopsy through circulating tumor DNA (ctDNA) analysis enables non-invasive detection of minimal residual disease (MRD) and early identification of cancer relapse, facilitating timely clinical intervention. However, detecting ctDNA in plasma samples with low tumor burden remains challenging due to the scarcity of mutant molecules, the background noise of sequencing errors and somatic mutations in normal cell-free DNA (cfDNA). Here, we present a mutation-informed fragmentomic framework and evaluate it on 90 stage III colorectal cancer patients with three years of follow-up. Using 712 serial whole-genome sequenced cfDNA samples (30×) with matched whole-genome sequencing of tumor tissue and germline DNA from buffycoat for each patient, we collected cfDNA fragments spanning tumor-derived somatic mutation positions and compared fragmentomic characteristics of mutation-bearing and non-mutated cfDNA fragments within the same sample. By leveraging fragment length and fragment end-motif patterns, our approach can distinguish cancer-positive from cancer-negative plasma samples without requiring model training or panel-of-normals calibration. The method achieved AUCs of 0.863 and 0.74 using fragment length and end motif features, respectively, and 0.871 when combined, outperforming tumor fraction estimates based on the frequency of mutated fragments (AUC=0.832). Integrating fragmentomic features with tumor fraction further improved performance, yielding an AUC of 0.873, indicating complementary signals between fragmentomic patterns and mutation burden. Aggregated analyses revealed ctDNA-specific patterns, including fragment shortening, motif enrichment of A/T ends, and depletion of C/G ends, directly linking fragmentomic features to tumor-derived cfDNA. Overall, mutation-informed fragmentomic profiling improves ctDNA detection beyond counting mutant reads and provides a scalable, training-free strategy for MRD assessment and early relapse detection while offering mechanistic insights into tumor-specific cfDNA biology.

## Introduction

Circulating tumor DNA (ctDNA), a fraction of cell-free DNA (cfDNA) originating from tumor cells, has emerged as a non-invasive and clinically accessible biomarker that offers important advantages over analyses based on tissue biopsies [1], [2]. ctDNA carries tumor-derived genetic and epigenetic alterations that reflect the molecular landscape of the underlying malignancy [3]. Owing to the short half-life of circulating DNA fragments, plasma ctDNA levels can reflect real-time tumor dynamics and have therefore been extensively explored for timely cancer detection and disease monitoring across multiple cancer types [4], [5], [6], [7]. In colorectal cancer, ctDNA analysis has demonstrated clinical utility for risk stratification, assessment of therapy efficacy, and detection of minimal residual disease [8], [9], [10], [11], [12], [13], [14].

Tumor burden can be quantified by detecting cancer-specific somatic mutations in plasma ctDNA [4], [15]. Tumor-informed strategies, which leverage matched tumor tissue to define patient-specific target mutations, can achieve high sensitivity for cancer detection and disease monitoring [12], [13], [16], [17]. However, mutation-based approaches are inherently challenged in early-stage cancer and minimal residual disease settings, where ctDNA abundance is extremely low and true variants are difficult to distinguish from sequencing artefacts [18], [19], [20]. As a result, reliance on somatic mutations alone may be insufficient for robust ctDNA detection, motivating the investigation of complementary ctDNA features.

Fragmentomic features of cfDNA, including but not limited to fragment size, end motifs, breakpoint motifs, and nucleosome occupancy, have been shown to differ systematically between cfDNA from cancer patients and healthy individuals [21], [22], [23], [24]. Tumor-derived fragments are generally shorter than non-tumor cfDNA [25], [26], [27], exhibit distinct end-motif patterns [28], [29], and reflect cancer-associated cleavage processes and altered nucleosome organization [30]. By integrating and modeling multiple fragment-level signals using machine learning or deep learning frameworks, prior studies have demonstrated improved performance in cfDNA-based cancer detection [23], [31]. Together, these findings highlight the potential of fragmentomic features to enhance ctDNA detection in oncology in addition to mutations.

Although fragmentomic features have shown strong potential for distinguishing cfDNA from cancer patients and healthy individuals, most existing studies derive these features from bulk cfDNA, analyzing all fragments collectively rather than focusing on tumor-derived fragments [22], [25]. Typically, the majority of cfDNA originates from normal hematopoietic cells and tumor-derived fragments constitute only a small fraction of the total cfDNA, especially in early-stage cancer or post-operative settings [32]. Tumor-specific mutations have been used to validate tumor-associated fragmentation properties, however, they are not typically used to define the fragment pool for fragmentomic analysis [23]. Consequently, bulk cfDNA fragmentomic patterns represent a mixture of tumor and non-tumor signals, which may obscure cancer-specific characteristics and reduce sensitivity in low tumor-burden contexts.

To overcome the dilution of tumor-specific signals in conventional fragmentomics analyses, we introduce a mutation-informed fragmentomics framework. We focus exclusively on cfDNA fragments spanning positions containing somatic mutations identified from matched tumor sequencing, thereby enriching ctDNA-derived signals. This strategy establishes a direct and interpretable connection between tumor-specific mutations and fragment-level characteristics. Importantly, it operates at the single-sample level and does not rely on supervised training, external reference cohorts, or fragment-level classification models. We applied this approach to 90 stage III colorectal cancer patients with three years of follow-up, comprising 712 serial whole-genome sequenced cfDNA samples, previously reported by [12]. By analyzing fragment length and end-motif features of mutation-spanning fragments, stratified by mutant-allele or reference-allele support, our approach enables both per-sample ctDNA classification and aggregated fragment-level analyses. Together, this study demonstrates that mutation-informed fragmentomics provides a biologically grounded and scalable approach for integrating fragmentomic features into cfDNA analysis, supporting robust ctDNA detection in low-tumor-burden settings while enabling mechanistic investigation of ctDNA-specific fragmentation patterns.

## Results

### Extraction of cfDNA Fragments Spanning Tumor-Specific Mutations

To determine whether fragments supporting tumor-specific mutations exhibit distinct fragmentomic features that improve sample-level detection, we analyzed cfDNA fragments spanning somatic mutation sites identified from matched tumor sequencing. Although these loci are enriched for ctDNA, they also contain background fragments from normal cfDNA. We therefore stratified mutation-spanning fragments into mutant-allele–supporting and reference-allele–supporting groups. We then evaluated whether fragmentomic features derived from these groups can distinguish ctDNA-positive from ctDNA-negative plasma samples. An overview of the study design and analytical workflow is shown in Figure 1.

**Figure 1.**
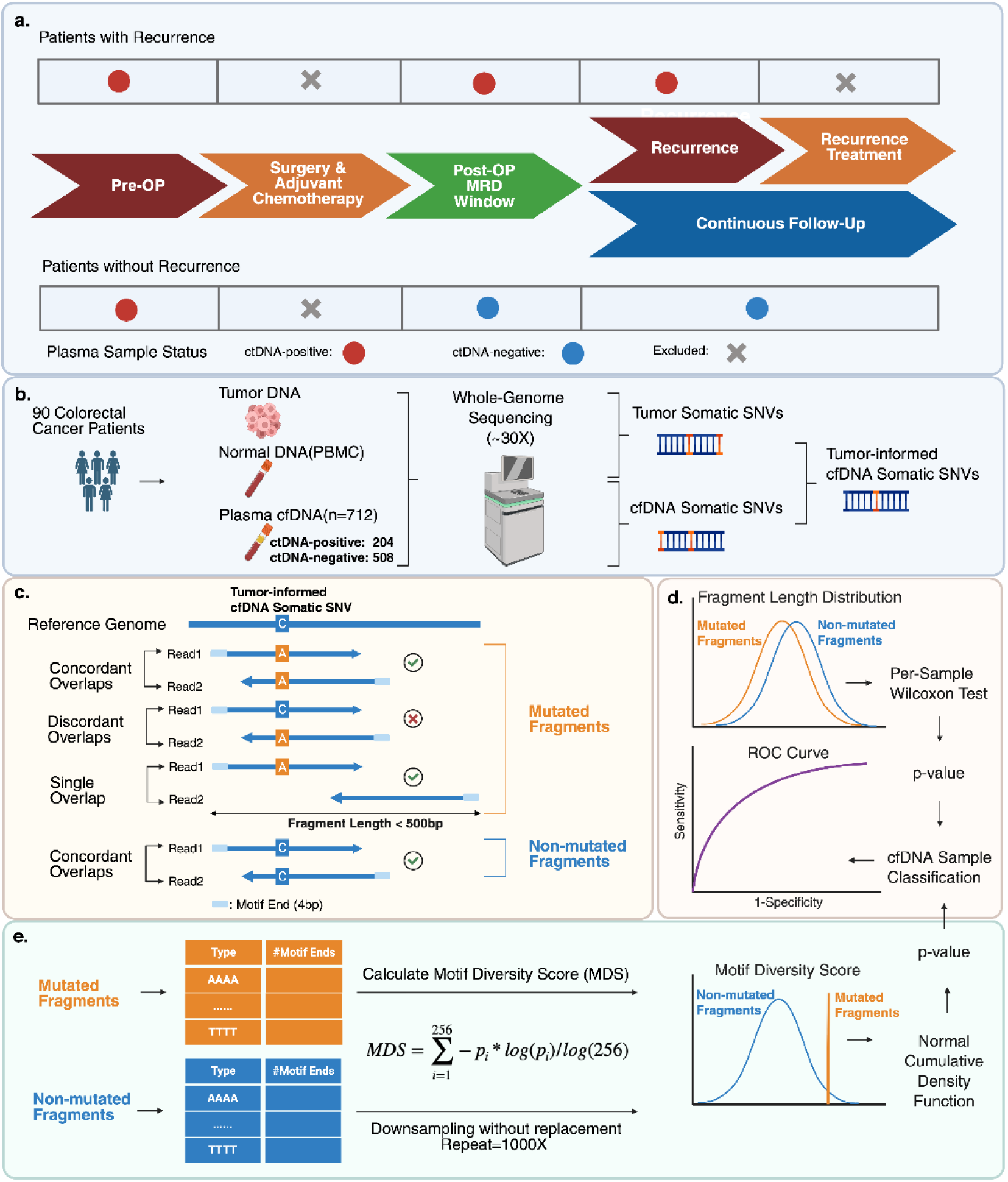
Systematic overview of the study design and analytical workflow. **a.** Time points of cfDNA sample collection relative to clinical interventions, including pre-operative, surgery and adjuvant chemotherapy (if administered), and post-operative MRD window. For patients who experienced relapse, additional time points included radiologically confirmed recurrence and recurrence-directed treatment; for non-relapsing patients, follow-up consisted of continued imaging surveillance. **b.** Available patient data and sequencing workflow. Whole-genome sequencing of tumor tissue, paired buffy coat, and cfDNA samples, followed by variant calling, resulted in the identification of tumor-informed cfDNA somatic SNVs. **c.** Extraction of mutation-spanning cfDNA fragments and their classification into mutated and non-mutated fragments. Fragment length and 5′-end 4-mer motif sequences were recorded as fragmentomic features. **d.** Fragment length–based sample classification.

We applied this mutation-informed fragmentomics framework to 90 stage III colorectal cancer patients with three years of follow-up, comprising 712 serial whole-genome sequenced cfDNA plasma samples. Among these, we labeled 204 samples as ctDNA-positive, including all pre-operative plasma samples and post-operative samples collected from patients with radiologically confirmed recurrence outside active treatment periods (Figure 1a). The remaining 508 ctDNA-negative samples were all post-operative plasma samples from non-recurrent patients with no radiological evidence of disease (Figure 1a).

Whole-genome sequencing of tumor tissue and matched buffy coat samples was used to identify somatic single nucleotide variants (SNVs) in the tumor. These SNVs were then intersected with somatic variants detected in cfDNA to define tumor-derived cfDNA mutations (Figure 1b) (details about the variant calling are provided in the methods section). cfDNA fragments spanning these mutation loci were extracted for downstream fragmentomic analyses. Fragments were then classified as mutated or non-mutated based on read-level allelic support at the mutation site. For each fragment, fragment length and 5′-end 4-mer motif composition were recorded as fragmentomic features (Figure 1c), forming the basis for subsequent analyses of mutation-informed fragment length and motif diversity.

A Wilcoxon rank-sum test was performed to compare fragment length distributions between mutated and non-mutated fragments within each cfDNA sample. The resulting p-values were used to rank samples and construct receiver operating characteristic (ROC) curves. **e.** Motif diversity score–based sample classification. Motif diversity scores were calculated from 5′-end motif frequencies of mutated fragments and from a randomly sampled equal-sized set of non-mutated fragments. This random sampling without replacement of non-mutated fragments was repeated 1000 times to get a distribution of non-mutated MDS scores. By comparing the motif diversity score of mutated fragments to the distribution derived from non-mutated fragments using a normal cumulative density function, p-values were obtained to rank samples and construct ROC curves.

### Mutation-Informed Fragmentomic Features Enable cfDNA Classification

#### Classification of cfDNA Samples Using Mutation-Informed Fragment Length

cfDNA fragments derived from tumor cells are known to be shorter than those originating from healthy tissues, with modal fragment lengths of approximately 150 bp and 167 bp, respectively. Based on this established observation, we hypothesized that in ctDNA-positive samples, mutated fragments would exhibit a length distribution shifted toward shorter fragments compared with non-mutated fragments at the same positions within the same cfDNA sample. To quantify the divergence between fragment length distributions on a per-sample basis, we applied a one-sided Wilcoxon rank-sum test, using the resulting p-value as a fragmentomic signal of ctDNA presence (Figure 1d).

To illustrate this approach, we compared a pre-operative cfDNA sample with a post-operative sample from the same non-relapsing patient (Figure 2a). In the post-operative sample, where no residual tumor was detected (tumor fraction of 0%, estimated by MRDetect), only a limited number of mutation-spanning fragments were identified, resulting in sparse fragment length profiles. No systematic shift was observed between the length distributions of mutated and non-mutated fragments. Non-mutated fragments exhibited two characteristic peaks, with a dominant peak around 170 bp and a secondary peak around 340 bp (Figure 2a). For statistical comparison, fragment lengths were restricted to the 80–220 bp range, within which the Wilcoxon test yielded a non-significant result (p-value = 0.164), consistent with classification as ctDNA-negative (Figure 2a).

**Figure 2.**
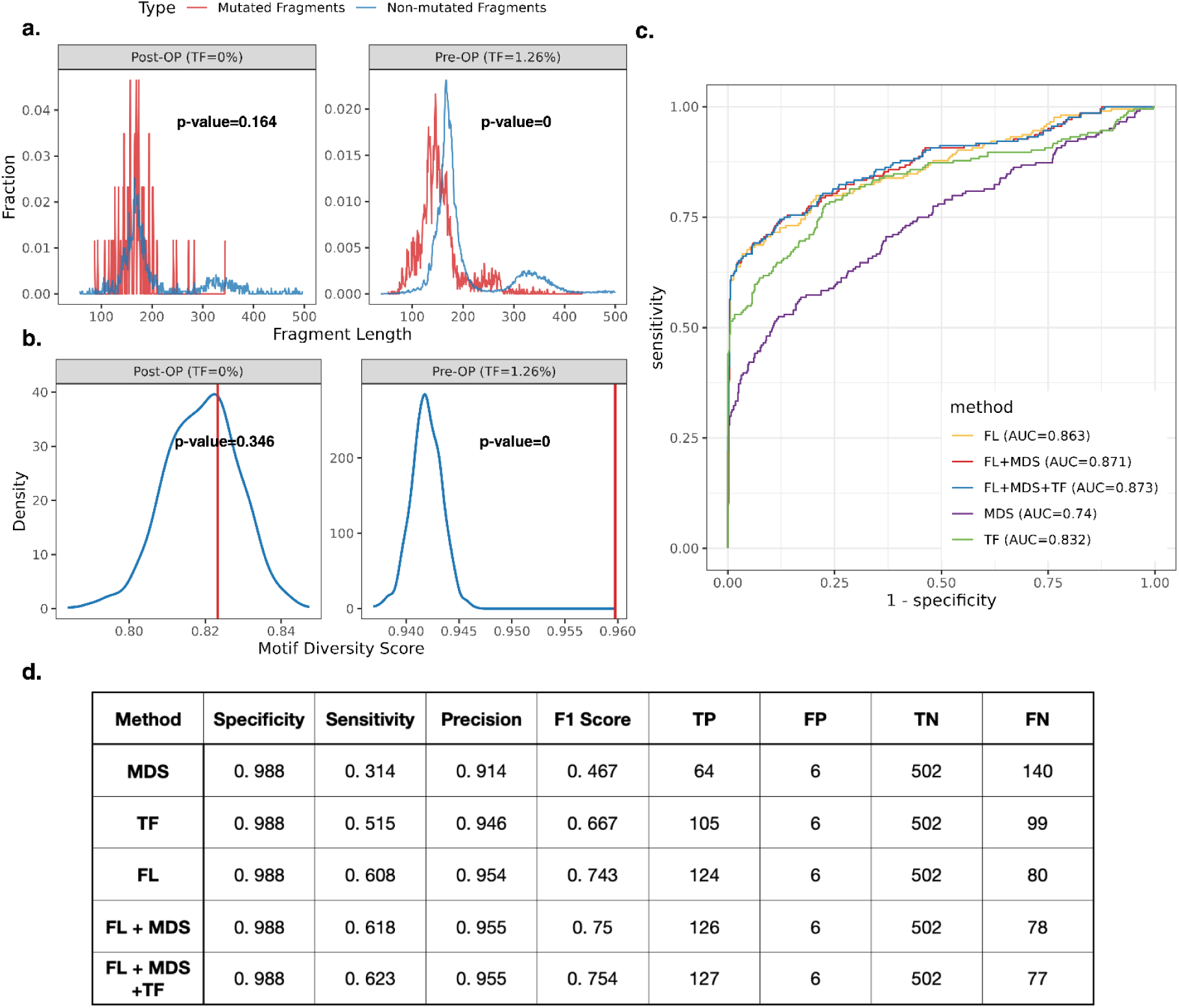
Classification of cfDNA samples using mutation-informed fragmentomic features. **a.** Fragment length distributions of mutated and non-mutated fragments for two representative cfDNA samples from the same patient: a post-operative sample with a tumor fraction of 0% and a pre-operative sample with a tumor fraction of 1.26%, both estimated by MRDetect. Statistical significance was assessed using the Wilcoxon rank-sum test. **b.** Motif diversity score (MDS) distributions for mutated and non-mutated fragments from the same representative samples are shown in panel a. Statistical significance was evaluated using a one-sided cumulative normal distribution. **c.** Receiver operating characteristic (ROC) curves comparing cfDNA sample classification performance using mutation-based tumor fraction (TF), fragment length (FL), motif diversity score (MDS), their combined fragmentomics model (FL+MDS), and the integrated model incorporating fragmentomics and tumor fraction (FL+MDS+TF). **d.** Summary performance metrics evaluated at a target specificity of 99%, reporting achieved specificity (98.8%), sensitivity, precision, F1 score, and confusion matrix counts, including true positive (TP), false positive (FP), true negative (TN), and false negative (FN).

In contrast, the pre-operative cfDNA sample (tumor fraction of 1.26%, estimated by MRDetect) showed a clear shift of mutated fragments toward shorter lengths. The primary peak of mutated fragments was centered around 150 bp, compared with approximately 170 bp for non-mutated fragments, and the secondary peak was shifted from ∼340 bp to ∼320 bp (Figure 2a). Accordingly, the Wilcoxon test (80–220 bp) produced a highly significant p-value (p-value = 0), indicating strong separation between mutated and non-mutated fragment length distributions and consistent with ctDNA positivity (Figure 2a).

#### Classification of cfDNA Samples Using Mutation-Informed End Motifs

We next evaluated whether mutation-informed end motifs provide an independent signal for cfDNA sample classification. Specifically, we extracted 4-mer sequences from the 5′ ends of cfDNA fragments. Previous studies have reported that end motif composition differs between cfDNA from healthy individuals and cancer patients [28]. To quantify this variation, we adopted the motif diversity score (MDS) in line with previous studies [28], which measures the uniformity of end motif composition within a sample. MDS ranges from 0 to 1, with higher values indicating a more diverse and more evenly distributed set of end motifs. Prior work has reported elevated MDS in cfDNA from hepatocellular carcinoma patients compared with healthy controls [28]. Based on these observations, we hypothesized that ctDNA-positive cfDNA samples would exhibit a larger difference in MDS between mutated and non-mutated fragments than ctDNA-negative samples.

Across all cfDNA samples, MDS exhibited a strong dependence on fragment count. Mutated fragments from ctDNA-positive samples showed higher MDS values than those from ctDNA-negative samples (Supplementary Figure 1a). However, non-mutated fragments consistently displayed substantially higher MDS values than mutated fragments in both groups, due to the higher number of fragments, obscuring the expected biological pattern in which cancer-derived cfDNA exhibits increased motif diversity relative to healthy cfDNA.

After normalizing fragment counts by sampling a set of non-mutated fragments that match the number of mutated fragments, MDS values for non-mutated fragments decreased to levels comparable with those of mutated fragments. Under this normalization, ctDNA-positive samples exhibited a larger median MDS difference between mutated and non-mutated fragments than ctDNA-negative samples, revealing a biologically consistent separation (Supplementary Figure 1b).

To quantify the statistical significance of motif diversity differences, we applied a one-sided cumulative normal distribution to assess whether the MDS of mutated fragments exceeded the empirical distribution of MDS values derived from non-mutated fragments (Figure 1e). Using the same paired cfDNA samples as in the fragment length analysis, we observed that mutated fragments exhibited higher MDS values than non-mutated fragments in each sample, with a pronounced shift toward higher MDS values in the ctDNA-positive sample relative to the ctDNA-negative sample (Figure 2b). The post-operative cfDNA sample with an estimated tumor fraction of 0% yielded a non-significant p-value of 0.346, indicating no meaningful elevation of MDS in mutated fragments (Figure 2b). In contrast, the pre-operative cfDNA sample with a tumor fraction of 1.26% showed a highly significant deviation (p-value = 0), consistent with ctDNA positivity (Figure 2b).

#### Benchmarking and Combined Performance

To benchmark sample-level ctDNA classification performance, we compared mutation-informed fragmentomic features with a mutation-only approach. For each cfDNA sample, we derived p-values from (i) fragment length differences between mutated and non-mutated fragments (FL), and (ii) motif diversity score differences (MDS). These were evaluated alongside a mutation-based tumor fraction (TF), defined as the proportion of fragments carrying the alternative allele among all fragments spanning tumor-informed somatic SNVs (Supplementary Figure 2a). TF served as a mutation-only baseline reflecting allelic burden without fragmentomic information. Classification performance was assessed using receiver operating characteristic (ROC) analysis and area under the curve (AUC).

As shown in Figure 2c, mutation-informed fragment length (FL) achieved the strongest classification performance individually, with an AUC of 0.863, outperforming the mutation-based tumor fraction (TF) (AUC = 0.832). This result indicates that fragmentomic features provide additional discriminative power beyond that provided by mutation information alone. The motif diversity score comparison (MDS) also enabled separation of samples labeled ctDNA-positive and ctDNA-negative, but yielded a lower AUC of 0.74.

Furthermore, combining fragment length and motif diversity score (FL+MDS) using Fisher’s method further improved classification performance, yielding an AUC of 0.871. To integrate mutation information robustly, we computed a TF-based p-value by comparing the observed tumor fraction at each sample’s real mutation sites to a set of randomly selected, nearby reference sites (≥200 bp away) and assessing the deviation using a cumulative normal distribution (Supplementary Figure 2b). This TF-based p-value was then combined with the fragmentomic features (FL+MDS) to produce a three-way integrated score (TF+FL+MDS), achieving the highest overall AUC of 0.873, although the improvement over FL+MDS was modest.

At 99% specificity, sensitivity increased from 51.5% for TF alone to 60.8% for FL, 61.8% for FL+MDS, and 62.3% for TF+FL+MDS (Figure 2d), reflecting a consistent reduction in false negatives while maintaining a fixed false-positive rate. These results demonstrate that p-value-based fragmentomic features comparisons capture most of the discriminative signal, while the addition of the background-corrected TF p-value yields a modest but consistent improvement in ctDNA classification.

We additionally compared our approach to MRDetect [16], a tumor-informed genome-wide SNV integration method that models tumor fraction as a function of mutation load and sequencing depth. At 99% specificity, MRDetect achieved a sensitivity of 73.5%, which is higher than that of our best combined model, TF+FL+MDS (62.3%) (Supplementary Figure 3). While mutation-count–based genome-wide integration remains highly sensitive, our framework provides an alternative strategy based on fragment-level biological features that does not rely on model training.

### Global Patterns of Aggregated Mutation-Spanning Fragments

Beyond per-sample classification, we explored global patterns of cfDNA fragmentomic features using mutation-spanning fragments. For proof-of-concept comparisons, we aggregated mutated fragments from ctDNA-positive cfDNA samples and non-mutated fragments from ctDNA-negative cfDNA samples, hereafter referred to as ctDNA-positive fragments and ctDNA-negative fragments, respectively. This aggregation yielded approximately 0.9 million ctDNA-positive fragments and 0.6 million ctDNA-negative fragments (Supplementary Figure 4). All downstream analyses were performed on these aggregated fragment sets.

We first examined whether fragment length distributions differed between the two aggregated fragment sets. Figure 3a shows that both sets share a major peak at ∼170 bp. For fragments shorter than 150 bp, periodic peaks at ∼10 bp intervals are apparent and particularly pronounced in mutated fragments from ctDNA-positive samples. In ctDNA-positive fragments, counts gradually decrease toward shorter lengths starting at ∼150 bp, whereas ctDNA-negative fragments exhibit only minor fluctuations. A secondary peak occurs around 300 bp for ctDNA-positive fragments, compared with ∼340 bp for ctDNA-negative fragments. These observations are consistent with known differences in fragment length distributions between cancer-derived and healthy cfDNA, supporting the validity of our aggregated fragment sets and the notion that mutation-spanning fragments capture cancer-associated cfDNA signals.

**Figure 3.**
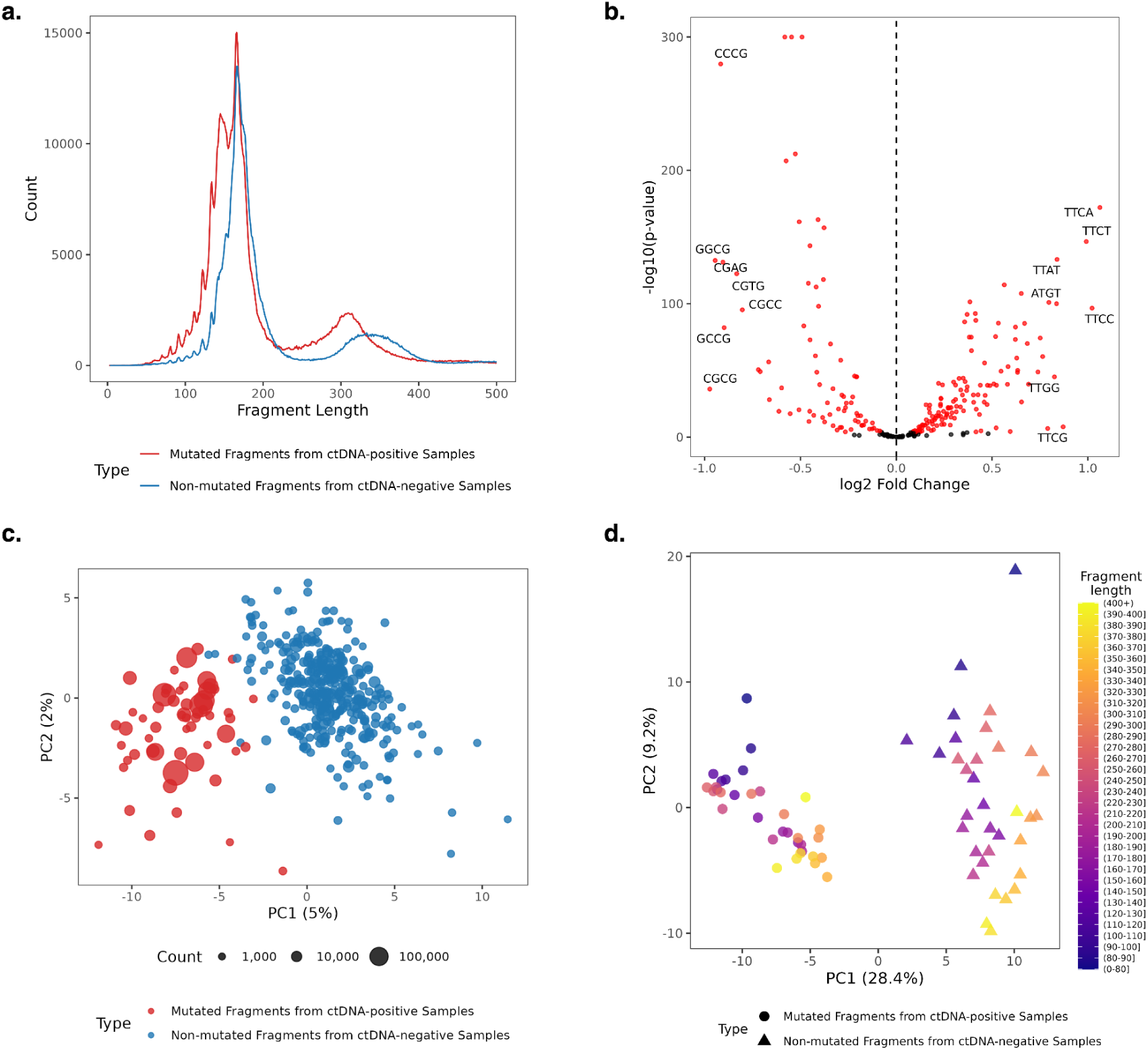
Global patterns of mutation-informed fragmentomics based on aggregated fragment sets. **a.** Fragment length distributions of aggregated mutated fragments from ctDNA-positive cfDNA samples (ctDNA-positive fragments) and non-mutated fragments from ctDNA-negative cfDNA samples (ctDNA-negative fragments). **b.** Volcano plot of 5′-end 4-mer motif composition comparing ctDNA-positive and ctDNA-negative fragment sets. Red points indicate motifs with Bonferroni-corrected *p-value*<0.05. Labeled motifs meet both significance (Bonferroni-corrected *p-value*<0.05) and effect size (|log₂ fold change|>0.8) thresholds. Positive log₂ fold change indicates the motif is more abundant in ctDNA-positive fragments compared to ctDNA-negative fragments, while negative log₂ fold change indicates it is less abundant. **c.** PCA plot of cfDNA samples retaining ≥1,000 fragments from the aggregated fragment set, based on end motif composition. **d.** PCA plot of ctDNA-positive and ctDNA-negative fragment sets, stratified by 10-bp length bins from 80 bp to 400 bp, with fragments <80 bp and >400 bp assigned to separate bins. Each group is represented by the proportions of all fragment end motifs.

We next analyzed end-motif enrichment and depletion of the two aggregated fragment sets. Comparing motif frequencies between ctDNA-positive and ctDNA-negative fragments (Figure 3b), we found that, while most end-motifs are statistically significant due to the large fragment counts, fold changes are generally modest. Using Bonferroni-corrected p-value<0.05 and |log2FC|>0.8 as thresholds, the top enriched motifs in ctDNA-positive fragments were TTCA, TTCC, TTCT, TTCG, TTAT, ATGT, and TTGG, and the top depleted motifs in ctDNA-positive fragments were CGCG, GGCG, CCCG, CGAG, GCCG, CGTG, and CGCC. Enriched motifs were predominantly T-end or A-end motifs, whereas depleted motifs were all C-end or G-end motifs, and all depleted motifs contained “CG”, likely reflecting methylation-driven differences in cleavage propensity at CpG sites [33]. These motif patterns further support the notion that mutation-spanning fragmentomics can detect ctDNA and inform cancer status.

To further explore whether end-motif composition could serve as a global biomarker, we performed principal component analysis (PCA) on the proportions of all 256 possible fragment end-motifs per sample calculated from aggregated fragments. Only cfDNA samples with at least 1,000 fragments remaining in the aggregated set were included, comprising 58 ctDNA-positive and 332 ctDNA-negative samples. Each point in Figure 3c represents a cfDNA sample characterized solely by fragments from the aggregated set. The first principal component (PC1) explains 5% of the variance and clearly separates ctDNA-positive and ctDNA-negative sample groups, with samples that intermingle generally having fewer fragments available. This analysis demonstrates that, beyond per-sample classification, mutation-spanning fragment end motifs exhibit consistent global patterns that distinguish ctDNA-positive from ctDNA-negative fragments, supporting the biological relevance of the per-sample features.

Finally, we investigated potential interactions between fragment length and end-motif composition, specifically whether fragments of similar lengths cluster together irrespective of their end-motifs. The aggregated fragments were grouped into 10-bp bins from 80 bp to 400 bp, with fragments shorter than 80 bp and longer than 400 bp assigned to separate bins at the lower and upper extremes, respectively. For each length bin, the proportions of all 256 possible fragment end motifs were calculated separately for mutated and non-mutated fragments, and these end motif profiles were used as input for PCA. As shown in Figure 3d, fragments within similar length bins showed a weak tendency to cluster along PC2, although this separation was limited and not pronounced. Only the shortest (<100 bp) and longest (>370 bp) fragment groups showed partial separation, and these groups still overlapped with neighboring length bins. In contrast, ctDNA-positive and ctDNA-negative samples remained clearly separated along PC1, driven primarily by end-motif composition. These results suggest that interactions between fragment length and end-motif features are modest, and that end-motif patterns represent a largely independent and dominant signal for distinguishing ctDNA status.

## Discussion

In this study, we demonstrate that cfDNA fragments spanning tumor-informed somatic mutations harbor distinct and biologically coherent ctDNA-associated fragmentomic patterns. Applied to a cohort of 90 stage III colorectal cancer patients, including 712 cfDNA samples collected over three years of follow-up, this mutation-informed fragmentomics strategy consistently improves cfDNA sample classification beyond mutation-based tumor fraction alone, highlighting the complementary diagnostic value of fragment length and fragment end-motif features that are directly linked to ctDNA presence. Beyond per-sample classification, aggregated analyses reveal ctDNA-specific fragmentomic signatures, including systematic fragment length shifts and end-motif biases, that are largely obscured in conventional fragmentomics analyses based on all cfDNA fragments. These findings establish mutation-informed fragmentomics as an effective and lightweight framework for enriching tumor-derived signals from a highly heterogeneous cfDNA background, while enabling biologically interpretable exploration of ctDNA-specific fragmentation patterns.

A key strength of the mutation-informed fragmentomics framework is its conceptual and practical simplicity. By restricting analyses to mutation-spanning fragments and leveraging direct within-sample comparisons between mutated and non-mutated fragments, the framework does not require supervised model training, parameter optimization, or calibration against external control cohorts. All fragmentomic features are computed directly from fragments within an individual cfDNA sample, enabling straightforward per-sample ctDNA assessment. The framework avoids cross-cohort generalization issues arising from batch effects that commonly affect machine learning models trained on fragmentomics data. This design reduces dependence on cohort composition and modeling assumptions while preserving a direct, interpretable link between somatic mutations and fragment-level characteristics. Therefore, mutation-informed fragmentomics provides a complementary and easily deployable strategy for integrating fragmentomic features into cfDNA analyses.

The effectiveness of mutation-informed fragmentomics likely arises from the coupling between tumor-specific genomic loci and altered DNA fragmentation processes in cancer cells. cfDNA fragmentation is influenced by nucleosome positioning, chromatin accessibility, and sequence-dependent nuclease cleavage, which differ systematically between tumor and normal cells due to epigenetic dysregulation and altered chromatin organization [23], [34]. By restricting fragmentomic analyses to cfDNA fragments spanning tumor-informed somatic mutations, our approach preferentially selects DNA molecules originating from tumor clones and situated within tumor-associated chromatin contexts [9], [35]. This locus-conditioned sampling allows direct comparison of mutant and reference fragments at tumor-specific genomic positions, enabling detection of ctDNA-specific fragmentation patterns, while substantially reducing background contributions from hematopoietic cfDNA. Consequently, it improves per-sample ctDNA classification sensitivity by jointly leveraging genomic information and fragmentomic features within a purified, mutation-spanning fragment pool.

Many of the fragmentomic patterns observed in our aggregated analyses, including shorter fragment lengths, 10-bp periodicity, and end-motif biases, are consistent with prior fragmentomics studies comparing bulk cfDNA from cancer patients and healthy individuals [23], [26], [30]. Notably, the enrichment of A or T end-motifs and the depletion of C or G end-motifs containing CpG dinucleotides observed in our colorectal cancer cohort have also been previously reported in hepatocellular carcinoma, supporting the notion that nuclease activity contributes similarly across cancer types [28]. Mechanistically, lower levels of DNASE1L3 in solid tissues and cancers relative to hematopoietic cells likely drive the depletion of 5′-CC tumor-fragments in ctDNA [28], [36], [37], [38], whereas DNASE1 and DFFB preferentially generate T-end and A-end fragments, respectively [37]. Importantly, our study demonstrates that these fragmentomic signatures are attributable to ctDNA, rather than reflecting population-level differences in bulk cfDNA.

Despite the demonstrated advantages, our study has several limitations. First, the mutation-informed fragmentomics framework relies on tumor-informed somatic mutations to enrich ctDNA, which requires matched tumor tissue and therefore limits its applications to settings where tumor tissue is available. Second, the statistical power of our method depends on the number of tumor-informed mutations and on their clonality. If the majority of the tumor-informed mutations are subclonal, a substantial fraction of the unmutated fragments will still be from the tumor, thus making the difference between mutated and unmutated fragments less clear. Finally, although our analyses included 712 cfDNA samples, the framework has only been evaluated in a colorectal cancer cohort, and its generalizability to other cancer types remains to be established.

In conclusion, we developed a mutation-informed fragmentomics approach that leverages cfDNA fragments that span tumor-informed somatic mutations to enrich ctDNA signals. Our method directly links genetic alterations with fragment-level features, improving per-sample ctDNA classification and revealing biologically consistent ctDNA-specific fragmentomic patterns. The framework is conceptually simple, requiring no external controls or supervised modeling, and can be applied to individual samples. Overall, it provides a scalable and interpretable strategy for minimally invasive cancer detection and for exploring ctDNA biology.

## Methods

### Sample Collection and Sequencing

This study included 90 patients diagnosed with UICC stage III colorectal cancer, recruited from seven Danish hospitals between November 2002 and February 2019. All patients received standard-of-care treatment and recurrence surveillance, including curative-intent tumor resection, adjuvant chemotherapy with 5-fluorouracil and oxaliplatin at the treating clinician’s discretion, and follow-up CT imaging at 12 and 36 months after surgery. Detailed patient information and associated clinical data are available in [12].

Plasma samples were collected before surgery, after surgery, and during follow-up visits every three months for up to three years. Based on longitudinal radiological imaging, which served as the clinical reference standard for ctDNA status, ctDNA-positive samples were defined as (i) all pre-operative cfDNA samples, and (ii) post-operative cfDNA samples from patients with radiologically confirmed recurrence, collected after completion of initial treatment (surgery with or without adjuvant chemotherapy) and before initiation of recurrence-directed therapy [39]. Conversely, ctDNA-negative samples were defined as post-operative cfDNA samples from patients without recurrence, collected after completion of all treatment and with no abnormalities detected on imaging [39]. According to these criteria, 712 cfDNA samples were obtained in total with known ctDNA status. 204 cfDNA samples were ctDNA-positive, while 508 were ctDNA-negative.

Tumor tissues were freshly frozen, and paired buffy coat samples were collected from each patient to enable filtering of germline and non-tumor variants. All tumor, buffy coat, and cfDNA samples underwent whole-genome sequencing (WGS) on the Illumina NovaSeq platform, using paired-end reads of 150 base pairs. cfDNA and buffy coat samples were sequenced to an average depth of 20×, while tumor samples were sequenced to a target coverage of 30× for those with histologically estimated tumor purity >30%, and 60× for samples with ≤30% purity. Detailed dataset information has been reported previously [12].

### Somatic SNV Detection

Sequencing data (FASTQ files) from tumor, buffy coat, and plasma samples were aligned to the human reference genome (hg38) using BWA-MEM [40]. The resulting BAM files were sorted and indexed with SAMtools [41]. Tumor and buffy coat BAM files were marked for duplicate reads using GATK [42] MarkDuplicates, followed by base quality score recalibration with GATK [42] BQSRPipelineSpark. The recalibrated BAM files were re-sorted and indexed with SAMtools [41].

Tumor somatic single nucleotide variants (SNVs) were called in tumor-normal mode using Mutect2 [43], with paired tumor and buffy coat BAM files. Variants were further filtered using GATK [42] FilterMutectCalls to remove non-PASS mutations and potential SNPs. The remaining high-confidence variants were defined as *Tumor Somatic SNVs* (Figure 1b) and used for downstream tumor-informed cfDNA mutation detection.

cfDNA somatic variant calling was performed using BBQ [44], which models sample-specific sequencing error rates and leverages discordant read overlaps to enable accurate mutation detection. During variant calling, bases within 5 bp of read ends were excluded, as were reads with low mapping quality, soft clipping, INDELs, or more than two mismatches relative to the reference genome. BAM files from the Buffy coat were used to filter out germline and non-tumor variants, and additional filtering removed variants present in gnomAD [45]. All remaining variants were considered as *cfDNA Somatic SNVs* (Figure 1b).

*Tumor-informed cfDNA Somatic SNVs* were subsequently defined as cfDNA somatic SNVs that overlapped with somatic SNVs identified in the matched tumor tissue.

### Mutation-Spanning Fragment Extraction

Fragments spanning *Tumor-informed cfDNA Somatic SNVs* were extracted for downstream cfDNA sample classification analyses. Fragment extraction was performed using pysam [46]. Fragments were retained only if they contained no INDELs or soft-clipped bases, had at most two mismatches relative to the reference, a minimum mapping quality of 50, and the mutation site was located at least 5 bp from either fragment end.

Fragments were then classified according to the allelic status at the mutation site. Fragments in which both reads carried the reference allele were defined as *non-mutated fragments*, whereas fragments with concordant mutant alleles were defined as *mutated fragments*. Discordant fragments, in which the two reads supported different alleles, were excluded. If only one read of a read pair overlapped the mutation site and the fragment length did not exceed 500 bp, the fragment was classified based on the allele observed in the overlapping read. (Figure 1c)

For each retained fragment, the fragment length and 5′-end 4-mer motif sequences were extracted for downstream analyses.

### CfDNA Sample Classification based on Fragmentomic Features Extracted from Mutation-Spanning Fragments

#### Fragment Length-based Sample Classification

Fragment length features were extracted from all qualified mutation-spanning cfDNA fragments, retaining only fragments with lengths between 80 and 220 bp. For each cfDNA sample, a one-sided Wilcoxon rank-sum test was performed in R [47] to compare the fragment length distributions of mutated and non-mutated fragments. We hypothesized that ctDNA-positive samples would exhibit greater divergence between the two distributions, as reflected by smaller p-values.

The resulting p-values were used to rank samples and to assess classification performance using receiver operating characteristic (ROC) curve analysis and the corresponding area under the curve (AUC), as implemented in the R pROC [48] package (Figure 1d).

#### Motif Diversity Score-based Sample Classification

Four-mer motif sequences were extracted from the 5′ ends of all qualified mutation-covering cfDNA fragments. For each cfDNA sample, the number of non-mutated fragments was downsampled to match the number of mutated fragments, sampling without replacement. This procedure was repeated 1000 times to account for sampling variability.

The motif diversity score (MDS) was calculated separately for mutated and non-mutated fragments using the formula 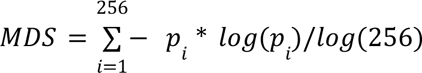 as described in [28], where *p_i_* denotes the frequency of the *i*-th 4-mer motif.

The empirical distribution of MDS values from non-mutated fragments was modeled using the cumulative distribution function (CDF) of a fitted normal distribution. We hypothesized that ctDNA-positive samples would exhibit a greater deviation of the MDS derived from mutated fragments toward the right tail of the non-mutated MDS distribution, reflected by smaller p-values.

The resulting p-values were used to rank samples and to assess classification performance using receiver operating characteristic (ROC) curve analysis and the corresponding area under the curve (AUC), as implemented in the R pROC [48] package (Figure 1e).

### Tumor Fraction

#### Tumor Fraction Calculation

Tumor fraction for each cfDNA sample was estimated as the ratio of the number of mutated cfDNA fragments overlapping *cfDNA Tumor-informed Somatic SNVs* to the sum of (i) the number of mutated cfDNA fragments overlapping *cfDNA Tumor-informed Somatic SNVs*, and (ii) the number of non-mutated cfDNA fragments overlapping all *Tumor Somatic SNVs* (Supplementary Figure 2a). This fraction reflects the proportion of cfDNA derived from tumor cells in the sample.

#### Tumor Fraction p-value Estimation Using Matched Reference Sites

To assess whether the observed tumor fraction at real mutation sites exceeded background noise, we computed a TF p-value using sequence-matched nearby reference sites to construct a local null distribution. For each somatic mutation, we generated 10 reference sites located at least 200 bp downstream of the real mutation position to avoid extracting the same mutated fragments from the true mutation sites (Supplementary Figure 2b). Starting from the +200 bp position, we sequentially identified the next 10 reference bases in the reference genome. Each reference site was assigned the same alternative allele as the corresponding real mutation to preserve nucleotide context and base-specific error characteristics. Consequently, 10 random mutations were created.

Tumor fraction was computed at both real and reference sites using the same criteria described above. The observed TF at the real mutation sites was then compared to the distribution of TF values across the 10 random mutation sites within the same sample. A TF p-value was calculated by evaluating the deviation of the observed TF from the reference distribution using a cumulative normal distribution.

### Benchmarking

For each cfDNA sample, p-values derived from the fragment length (FL) and motif diversity score (MDS) analyses were compared with the corresponding tumor fraction (TF) by generating receiver operating characteristic (ROC) curves and calculating the area under the curve (AUC). Additionally, the p-values from FL and MDS were combined using Fisher’s method (FL+MDS) to evaluate the performance of the integrated fragmentomic signal.

An additional combined metric incorporating fragmentomic features and the background-controlled TF p-value (FL+MDS+TF) was also evaluated. In total, five methods were benchmarked.

Performance was further compared to tumor fraction estimates generated by MRDetect [16], providing an external reference for performance evaluation.

### Exploratory Analyses Based on Aggregated Fragments

To investigate fragment length and end motif features at a global level, cfDNA data were aggregated from the sample level to the fragment level. Two fragment sets were generated: all mutated fragments from ctDNA-positive samples were pooled to form the ctDNA-positive fragment set, and all non-mutated fragments from ctDNA-negative samples were pooled to form the ctDNA-negative fragment set. This resulted in 907,280 ctDNA-positive fragments and 651,177 ctDNA-negative fragments (Supplementary Figure 4), which were used for subsequent exploratory analyses, including a comparison of fragment length distributions, motif enrichment analysis, and principal component analysis (PCA).

Fragment length distributions were visualized using ggplot2 in R, based on fragment counts at each base-pair length. For motif enrichment analysis, volcano plots were used to visualize the *log*_2_ fold change (*log*_2_ *FC*) versus − *log*_10_ (*p*–*value*) for each end motif, comparing ctDNA-positive fragments with ctDNA-negative fragments.

To assess whether end motif composition could serve as a potential biomarker for distinguishing ctDNA-positive and ctDNA-negative samples as a proof of concept, PCA was performed using the R [47] function prcomp on 4-mer end motif proportions derived from cfDNA samples with more than 1,000 fragments in the aggregated fragment set. This analysis included 58 ctDNA-positive and 332 ctDNA-negative cfDNA samples.

To further investigate potential interactions between end motif composition and fragment length, the same set of samples was stratified by fragment length into bins of 10 bp, spanning 80-400 bp, with fragments shorter than 80 bp and longer than 400 bp grouped separately. PCA was then performed again using R’s prcomp function to evaluate whether samples from shorter fragment length ranges exhibited distinct clustering patterns. In this analysis, each point represents a sample-specific fragment length bin.

## Code Availability

The workflows and analytical code used in this study are available at https://github.com/BesenbacherLab/Mutation_Informed_Fragmentomics.

## Data Availability

The data analysed in this article has previously been published in [12]. To protect the privacy and confidentiality of patients in this study, personal data, including clinical and sequence data, are not made publicly available in a repository or the supplementary material of the article. The data can be requested at any time from the corresponding authors. Any requests will be reviewed within a time frame of 2 to 3 weeks by the data assessment committee to verify whether the request is subject to any intellectual property or confidentiality obligations. All data shared will be de-identified. Request for access to raw sequencing data furthermore requires that the purpose of the data re-analysis is approved by The Danish National Committee on Health Research Ethics. The clinical and WGS data used in the study are available through controlled access from GenomeDK (https://genome.au.dk/library/GDK000005/).

## Author contributions

Conceptualization, Y.L., C.L.A. and S.B.; Methodology, Y.L., C.L.A. and S.B.; Software, Y.L.; Validation, Y.L., Formal analysis, Y.L.; Investigation, Y.L.; Resources, A.F., M.H.R., C.L.A.; Data curation, Y.L., C.O., A.F., M.H.R.; Writing–original draft preparation, Y.L.; Writing–review and editing, Y.L., C.L.A. and S.B.; Visualization, Y.L.; Supervision, C.L.A. and S.B.; Project administration, Y.L., C.L.A. and S.B.; Funding acquisition, C.L.A. and S.B.

All authors have read and agreed to the published version of the manuscript.

## Acknowledgements

This work was supported by the Novo Nordisk Foundation (grant no. NNF21OC0069056 to S.B. and grant no. NNF22OC0074415 to C.L.A.), and the Danish Cancer Society (grant numbers R352-A20664 (C.L.A.), R231-A13845 (C.L.A.), and R257-A14700 (C.L.A.)).

## Supplementary

**Supplementary Figure 1.**
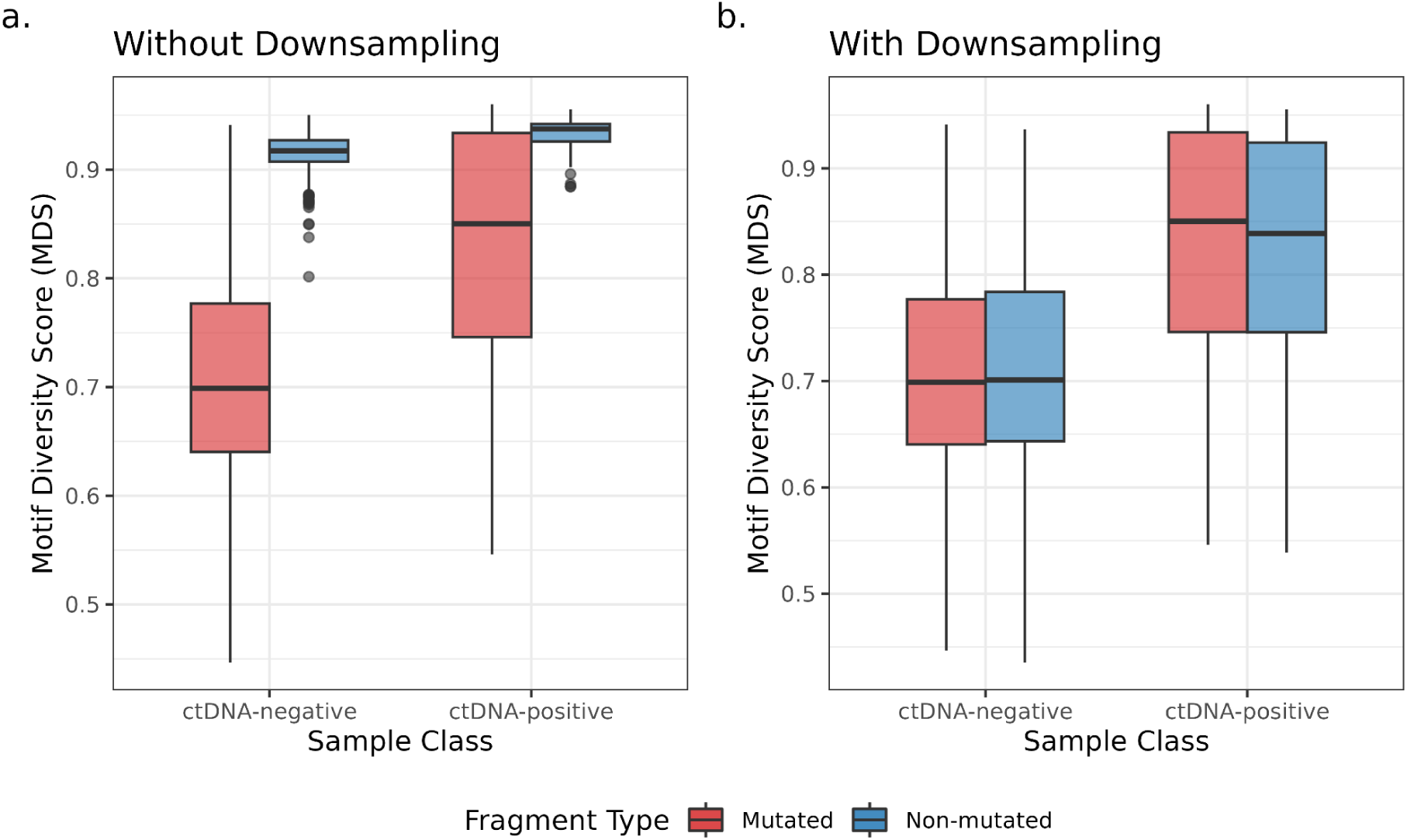
Distribution of Motif Diversity Scores (MDS) for mutated and non-mutated fragments in ctDNA-negative and ctDNA-positive cfDNA samples. **a.** Boxplot of raw MDS comparing mutated and non-mutated fragments for ctDNA-positive and ctDNA-negative samples. **b.** Boxplot of MDS comparing mutated fragments with the mean MDS of 100 downsampling iterations of non-mutated fragments for ctDNA-positive and ctDNA-negative samples.

**Supplementary Figure 2.**
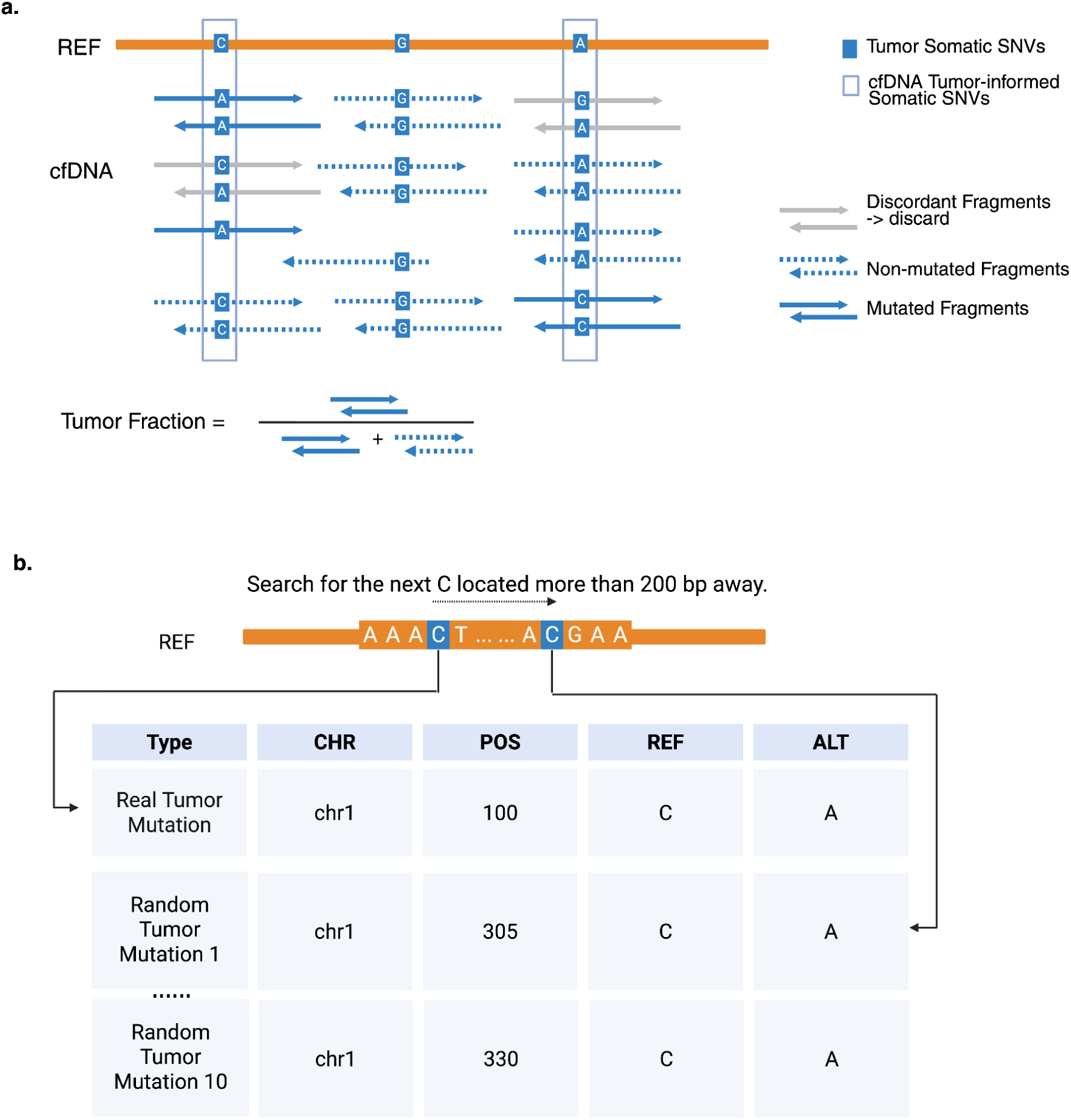
Overview of the method for estimating mutation-based tumor fraction. **a.** Tumor fraction is defined as the proportion of mutated fragments overlapping tumor-informed cfDNA somatic mutations relative to the sum of these fragments and all non-mutated fragments overlapping tumor somatic mutations. **b.** For each real mutation site, 10 nearby sequence-matched reference sites were generated by scanning ≥200 bp downstream and selecting the next 10 reference bases. Each reference site was assigned the same alternative allele as the real mutation. The tumor fraction at the real mutations was compared with the distribution across the random mutations to compute a TF p-value.

**Supplementary Figure 3.**
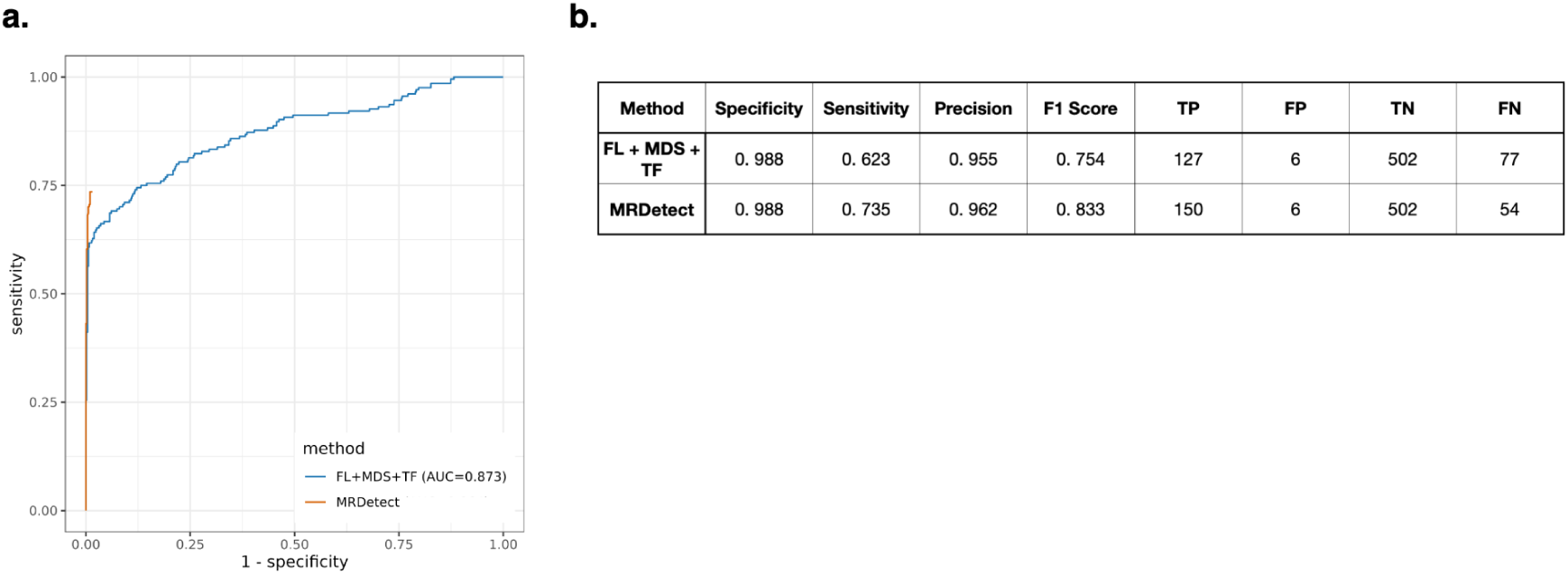
Comparison of cfDNA sample classification performance between the integrated method (FL+MDS+TF) and MRDetect. **a.** Receiver operating characteristic (ROC) curves comparing FL+MDS+TF and MRDetect. MRDetect has a reported lower limit of detection of TF= 10^−6^. **b.** Summary performance metrics evaluated at a target specificity of 99%, reporting achieved specificity (98.8%), sensitivity, precision, F1 score, and confusion matrix counts including true positive (TP), false positive (FP), true negative (TN), false negative (FN).

**Supplementary Figure 4.**
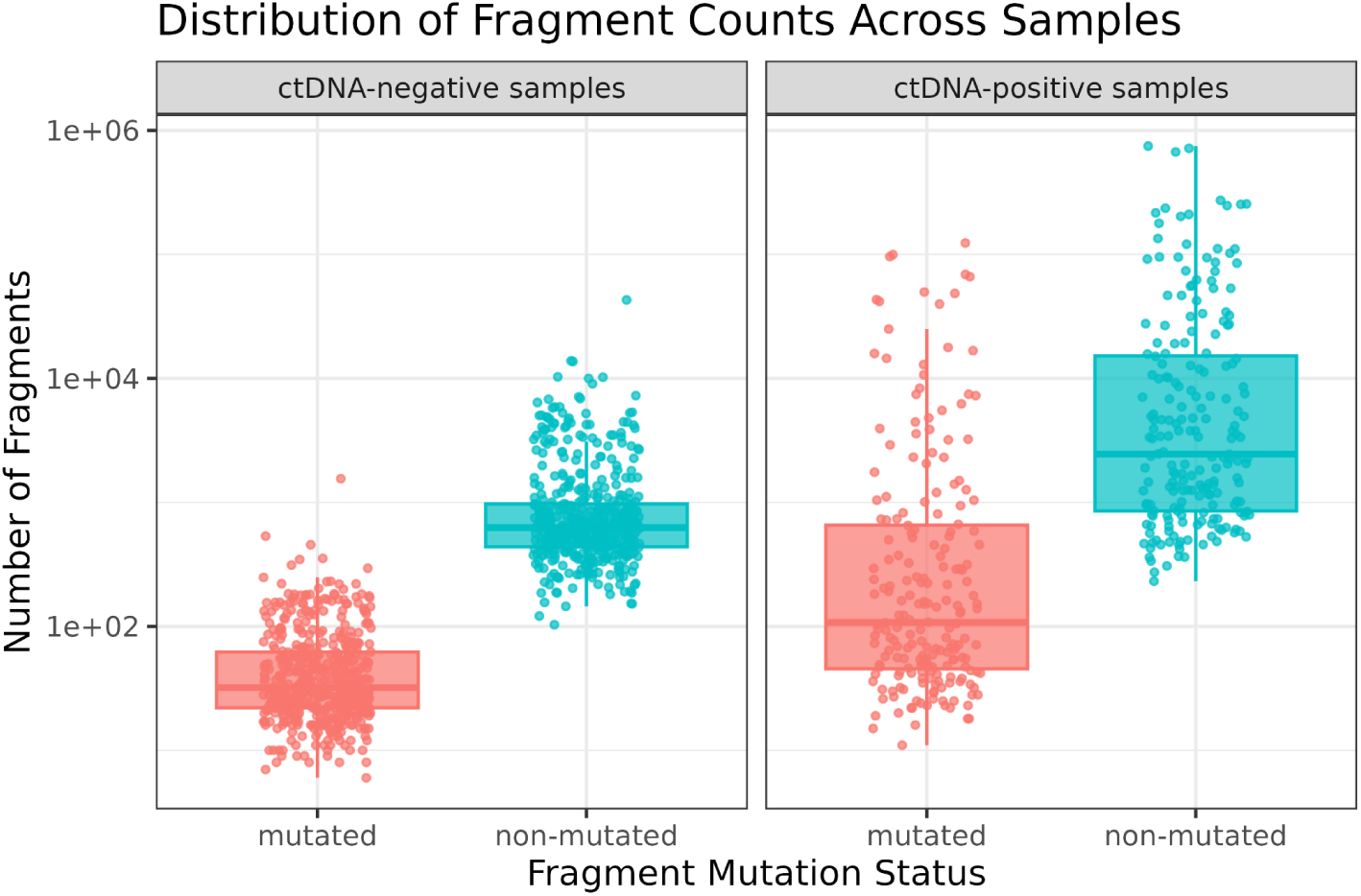
Distribution of fragment counts across cfDNA samples. Boxplots show the number of fragments stratified by mutation status (mutated vs. non-mutated) and sample type (ctDNA-positive vs. ctDNA-negative). In total, ctDNA-positive samples contain 907,280 mutated fragments and 6,820,738 non-mutated fragments, while ctDNA-negative samples contain 29,669 mutated fragments and 651,177 non-mutated fragments.

